# Early detection of late blight in potato by whole-plant redox imaging

**DOI:** 10.1101/2022.09.04.506513

**Authors:** Matanel Hipsch, Yaron Michael, Nardy Lampl, Omer Sapir, Yigal Cohen, Helman David, Shilo Rosenwasser

## Abstract

Late blight caused by the oomycete *Phytophthora infestans* is a most devastating disease of potatoes (*Solanum tuberosum)*. Its early detection is crucial for suppressing disease spread. Necrotic lesions are normally seen in leaves at 4 dpi (days post inoculation) when colonized cells are dead, but early detection of the initial biotrophic growth stage, when the pathogen feeds on living cells, is challenging. Here, the biotrophic growth phase of *P. infestans* was detected by whole-plant redox imaging of potato plants expressing chloroplast-targeted reduction-oxidation sensitive green fluorescent protein (chl-roGFP2). Clear spots on potato leaves with a lower chl-roGFP2 oxidation state were detected as early as 2 dpi, before any visual symptoms were recorded. These spots were particularly evident during light-to-dark transitions and reflected mislocalization of chl-roGFP2 outside the chloroplasts, demonstrating perturbation of the chloroplast import system by the pathogen. Image analysis based on machine learning enabled systematic identification and quantification of spots and unbiased classification of infected and uninfected leaves in inoculated plants. Comparing redox to chlorophyll fluorescence imaging showed that infected leaf areas which exhibit mislocalized chl-roGFP2 also showed reduced non-photochemical quenching (NPQ) and enhanced quantum PSII yield (ΦPSII) compared to the surrounding leaf areas. The data suggest that mislocalization of chloroplast-targeted proteins is an efficient marker of late blight infection and demonstrate how it can be utilized for nondestructive monitoring of the disease biotrophic stage using whole-plant redox imaging.

## Introduction

As the third major food crop, potato (*Solanum tuberosum*) is crucial for food security worldwide, providing starch, proteins, vitamins, and minerals (Devaux et al., 2014). Globally, its importance is growing rapidly, as manifested by the dramatic increase in potato production and demand in Asia, Africa, and Latin America over the past 20 years (da FAOSTAT, 2014). Despite the adaptability of potato to a wide range of environmental and climatic conditions, its productivity is highly affected by abiotic and biotic stresses (Bohnert, 2007). Specifically, late blight disease, which gave rise to the Irish Potato Famine, causes global annual losses of ∼6 billion dollars due to control expenditures and product damage (Haverkort et al., 2009). The causal agent of the disease is the oomycete plant pathogen *Phytophthora infestans* (Mont. DeBary). It can rapidly destroy leaves and tubers, and its extremely high reproductive potential renders it one of the most challenging pathogens to manage (Khan et al., 2017). *P. infestans* is a hemibiotrophic pathogen. It suppresses the immune system of the host during the early infection phase and spreads throughout the plant tissue while feeding on living cells without producing visual symptoms. Necrotic lesions, signifying the pathogen lifestyle switch to induced cell death and utilization of the nutrients from the dead cells, typically occur several days post-infection, leading to the spread of the diseases due to the massive sporangia production (Abramovitch & Martin, 2004; Cohen et al., 1997; Fry, 2008; Grenville-Briggs et al., 2005; Kamoun et al., 1997; Kanneganti et al., 2006; Kelley et al., 2010; Oh et al., 2009; Rubin et al., 2001; Tian et al., 2004; Van Damme et al., 2012, 2012). To better explore the molecular mechanism underlined this host-pathogen interaction and minimize crop losses and reduce fungicides inputs in agricultural practices, new technologies are required for early detection of the cellular events induced by the pathogen before the onset of symptoms.

Increasing evidence points to the role of chloroplast in sensing and responding to pathogens. Specifically, many chloroplast-localized effector proteins targeting chloroplast processes have been identified (de Torres Zabala et al., 2015; Gao et al., 2020; Kretschmer et al., 2020; Littlejohn et al., 2021; Tang et al., 2020; Xu et al., 2019). For example, it has been demonstrated that the interaction between AVRvnt1, a host-translocated RxLR type of effector protein, and chloroplast-targeted glycerate 3-kinase triggers a light-dependent immune response to *P. infestans* infection (Gao et al., 2020). The chloroplast, a central hub for plant metabolism, plays a central role in plant immune responses by metabolizing several key defense-related molecules, such as hormones, secondary metabolites, antioxidants and reactive oxygen species (ROS) (de Torres Zabala et al., 2015; Lee & Hwang, 2020; Serrano et al., 2016). Specifically, the overproduction of photosynthesis-derived ROS was shown to play a pivotal role in pathogen-associated molecular patterns (PAMPs) and effector-triggered immunity by its activation of the hypersensitive response (HR) and reprogramming of the transcription of genes involved in response to pathogens (Ding et al., 2019; Littlejohn et al., 2021). Accordingly, increasing ROS levels by silencing chloroplast-targeted ascorbate peroxidase (tAPX) or hampered glutathione biosynthesis, affected the expression of a large set of genes involved in response to pathogens (Ball et al., 2004; Maruta et al., 2012). Oxygen photoreduction at PSI during the water-water cycle (WWC) (Mehler reaction) (Mehler, 1951) is the primary source of ROS in the chloroplast. In addition, the interaction between molecular oxygen and the excited triplet state of chlorophyll at PSII produces singlet oxygen (Dogra et al., 2019; Mullineaux et al., 2018), a highly reactive molecule involved in lipid peroxidation (Asada, 1999; Kretschmer et al., 2020; Trotta et al., 2014). It has been suggested that H_2_O_2_ derived from oxygen photoreduction at PSI is involved in PAMP-triggered immunity, while rapid accumulation of singlet oxygen at PSII during effector-triggered immunity elicits PSII breakdown and HR (Littlejohn et al., 2021). In addition, chloroplast clustering around the nucleus, a phenomenon associated with a wide range of plant diseases, was shown to be mediated by H_2_O_2_ production (Ding et al., 2019).

Detoxification of chloroplastic H_2_O_2_ to water is mainly mediated by ascorbate peroxidases and peroxiredoxins (Awad et al., 2015). As electrons from reduced glutathione (GSH) are channeled to regenerate oxidized ascorbate via the ascorbate–glutathione pathway, H_2_O_2_ reduction gives rise to oxidized glutathione (GSSG) accumulation in plants (Foyer & Noctor, 2011;Rahantaniaina et al., 2013). High spatiotemporal-resolution-monitoring of glutathione redox potential (*E*_*GSH*_) can be achieved using redox-sensitive green fluorescent proteins (roGFPs) (Meyer et al., 2007; Meyer & Dick, 2010), which carry two engineered cysteine residues that form an intramolecular disulfide bridge which impacts its fluorescence characteristics (Dooley et al., 2004; Hanson et al., 2004). roGFP-based biosensors are effective tools for exploring redox dynamics in subcellular compartments at high spatiotemporal resolution in plants (Bratt et al., 2016; Gutscher et al., 2008; Jiang et al., 2006; Lampl et al., 2022; Meyer, 2008; Meyer & Dick, 2010; Nietzel et al., 2019; Schwarzländer et al., 2016; Ugalde et al., 2020, 2022). Spatially resolved mapping of stress-induced long-term redox perturbations has recently been demonstrated by whole-plant imaging of potato plants expressing roGFP2 in their chloroplasts (Hipsch et al., 2021b).

The present work employed a chloroplast-targeted roGFP (chl-roGFP2) sensor to examine *in planta E*_*GSH*_ dynamics in response to *P. infestans* infection. Leaf spots displayed a highly oxidized roGFP2 state were associated with the necrotrophic stages in the life cycle of the pathogen. Interestingly, we detected mislocalization of chl-roGFP2 outside the chloroplast during the biotrophic phase of the disease.That mislocalization affected the biosensor redox state upon light-to-dark transition and enabled detection of early infection by using whole-plant redox imaging. roGFP2 mislocalization was also associated with alterations in PSII efficiency and non-photochemical quenching (NPQ) activation, as demonstrated by chlorophyll fluorescence imaging (CFI). The results may have important implications on the detection of the early stages of *P. infestans* development, on the whole plant level, in potato plants under controlled and field conditions.

## Results

### *P. infestans* induces patches with an altered redox state on potato leaves in the necrotrophic stage of infection

We sought to investigate whether the chloroplastic *E*_*GSH*_ is altered during infection of potato leaves by *P. infestans* by monitoring the redox state of the chloroplast-targeted roGFP2 biosensor (chl-roGFP2), which allows real-time readout of *E*_*GSH*_ in living cells (Albrecht et al., 2011; Meyer et al., 2007). As shown in Fig 1. A-B, whole-leaf ratiometric images generated by dividing pixel by pixel images acquired following 405nm or 465nm excitations (R_405/465_) captured the changes in chl-roGFP2 in response to exogenous treatment with H_2_O_2_ and Dithiothreitol (DTT). R_405/465_ values of 0.084 and 0.42 were recorded in detached leaves treated with DTT and H_2_O_2,_ respectively, demonstrating a dynamic range of 5.1 (R_405/465_ for fully oxidized state divided by R_405/465_ for fully reduced state). These results validated the sensitivity of chl-roGFP2 to redox alterations and demonstrated the ability to spatially resolve these changes using whole-leaf fluorescence imaging.

**Figure 1:**
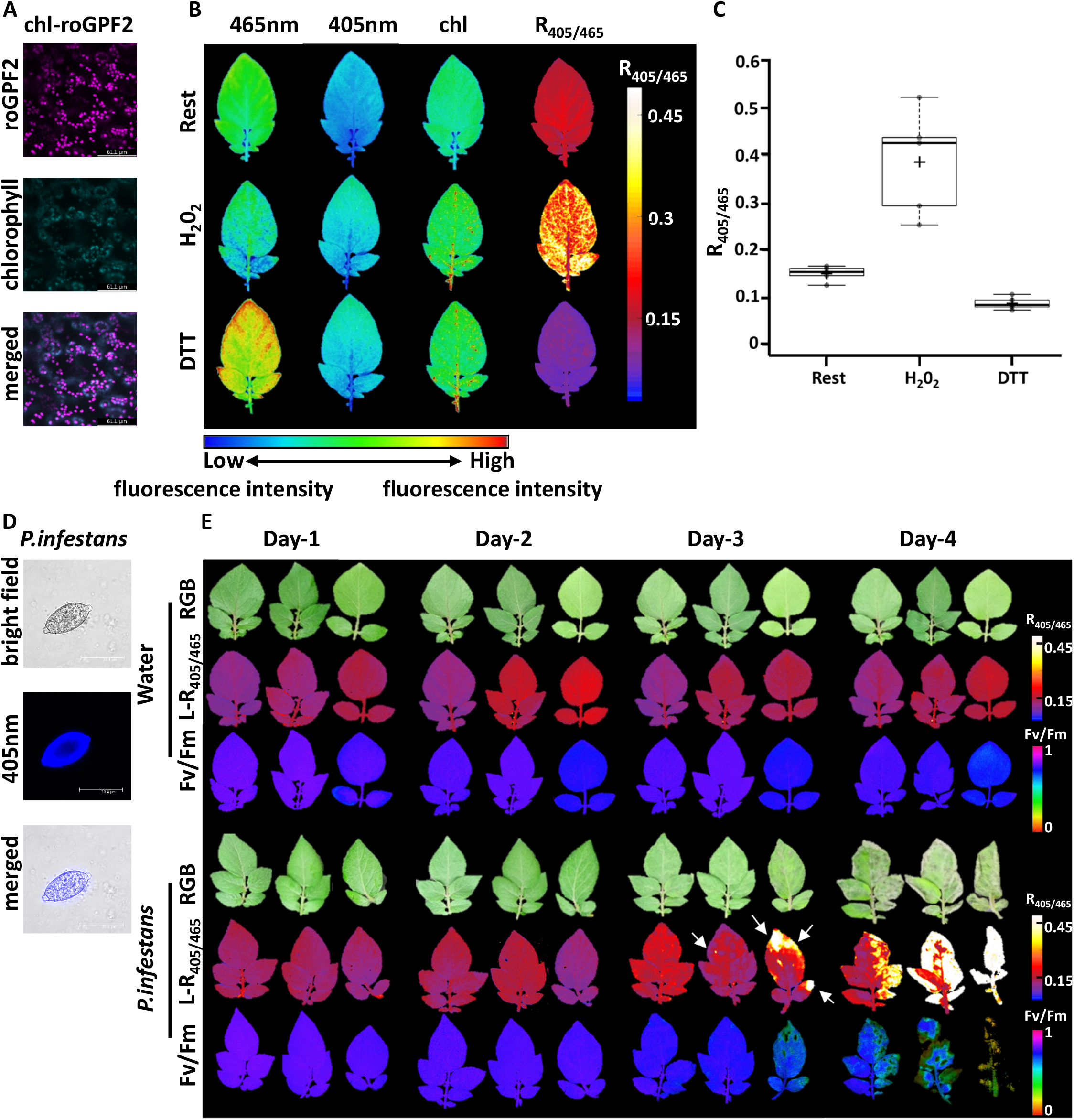
Spatially resolved redox changes in detached potato leaves during *P. infestans* infection. A) Confocal images showing the fluorescence signals emanating from chl-roGFP2-expressing potato plants (chl-roGFP2 excitation: 488 nm). B) Fluorescence signals measured following 405 and 465 nm excitation in detached leaves and ratiometric analysis of chl-roGFP2 signals during rest and under fully oxidized (1,000 mM H_2_O_2_) and fully reduced (100 mM DTT) conditions. Images of detached leaves were digitally combined for comparison. C) Quantification of Redox images presented in A. Values represent mean ± SE (n□= □5-7). Centerlines show the medians; box limits indicate the 25th and 75th percentiles; whiskers extend 1.5 times the interquartile range from the 25th and 75th percentiles. D) Confocal microscopy images of *P. infestans* stained with calcofluor-white (excitation: 405 nm). E) RGB, redox images of light-adapted potato leaves (L-R_405/465_) and maximum quantum yield of PSII (Fv/Fm).

To examine the effect of *P. infestans* on the chl-roGFP2 redox state, detached potato leaves of WT (wild type) and chl-roGFP2-expressing potato lines were inoculated by spraying with freshly harvested sporangia of *P. infestans* (Fig 1. D) or with distilled water (DW) as control. chl-roGFP2 ratiometric images were acquired from light-adapted leaves (L-R_405/465_) 3 h after light onset for five consecutive days. L-R_405/465_ measurements were compared to RGB images and images of the maximum quantum yield of PSII (Fv/Fm), a chlorophyll fluorescence-based indicator of plant physiological stress.

As shown in Fig 1. E, apparent symptoms of late blight, manifested as circular, necrotic brown and black spots surrounded by chlorotic tissue, were visible through RGB imaging, starting on the third day post-inoculation (dpi), and became more noticeable at 4 dpi. Sporangia of *P. infestans* were produced on the surface of the infected leaf tissues, the last stage in the asexual life cycle of this oomycete (Hardham, 2007). No signs of infection were observed on the control plants during the experiment. Interestingly, L-R_405/465_ images showed leaves with distinct oxidized or reduced spots at 3 and 4 dpi. At this stage, patches with lower Fv/Fm values were detected in the infected leaves (Fig 1, E & Fig, 1 supp). Spots showing a highly oxidized chl-roGFP2 state were generally associated with visible disease symptoms, while spots showing a reduced chl-roGFP2 state were symptomless. The magnitude of changes in the chl-roGFP2 oxidation state was comparable with previously recorded data collected from potato plants subjected to various abiotic stresses (Hipsch et al., 2021, Supp. Fig. 2). The results indicate that increased oxidation of roGFP2 is primarily associated with the necrotrophic stage of the late blight disease.

### Mislocalization of chl-roGFP2 outside the chloroplast is associated with the presymptomatic stages of *P. infestans* infection

To better understand the cellular events associated with the observed changes, the fluorescence signals emitted from roGFP2 were visualized by confocal microscopy at 2 and 4 dpi, representing the biotrophic and necrotrophic phases of the disease, respectively (Fig. 2). Surprisingly, while chl-roGFP2 was localized to chloroplasts in control leaves (Fig. 2 A-D), as reflected by the colocalization of GFP and chlorophyll autofluorescence signals, it was mislocalized in cells of infected leaf areas. In such tissue, GFP-related signals did not colocalize with chlorophyll autofluorescence. This chl-roGFP2 mislocalization was accompanied by chloroplast aggregation, which was seen as early as 2 dpi, before the onset of cell death or *P. infestans* sporulation (Fig. 2 and Supp. Fig 3&4). These observations imply the inhibition of chloroplast protein import during *P. infestans* infection and suggest that the redox changes in chl-roGFP2 oxidation are related to its mislocalization.

**Figure 2:**
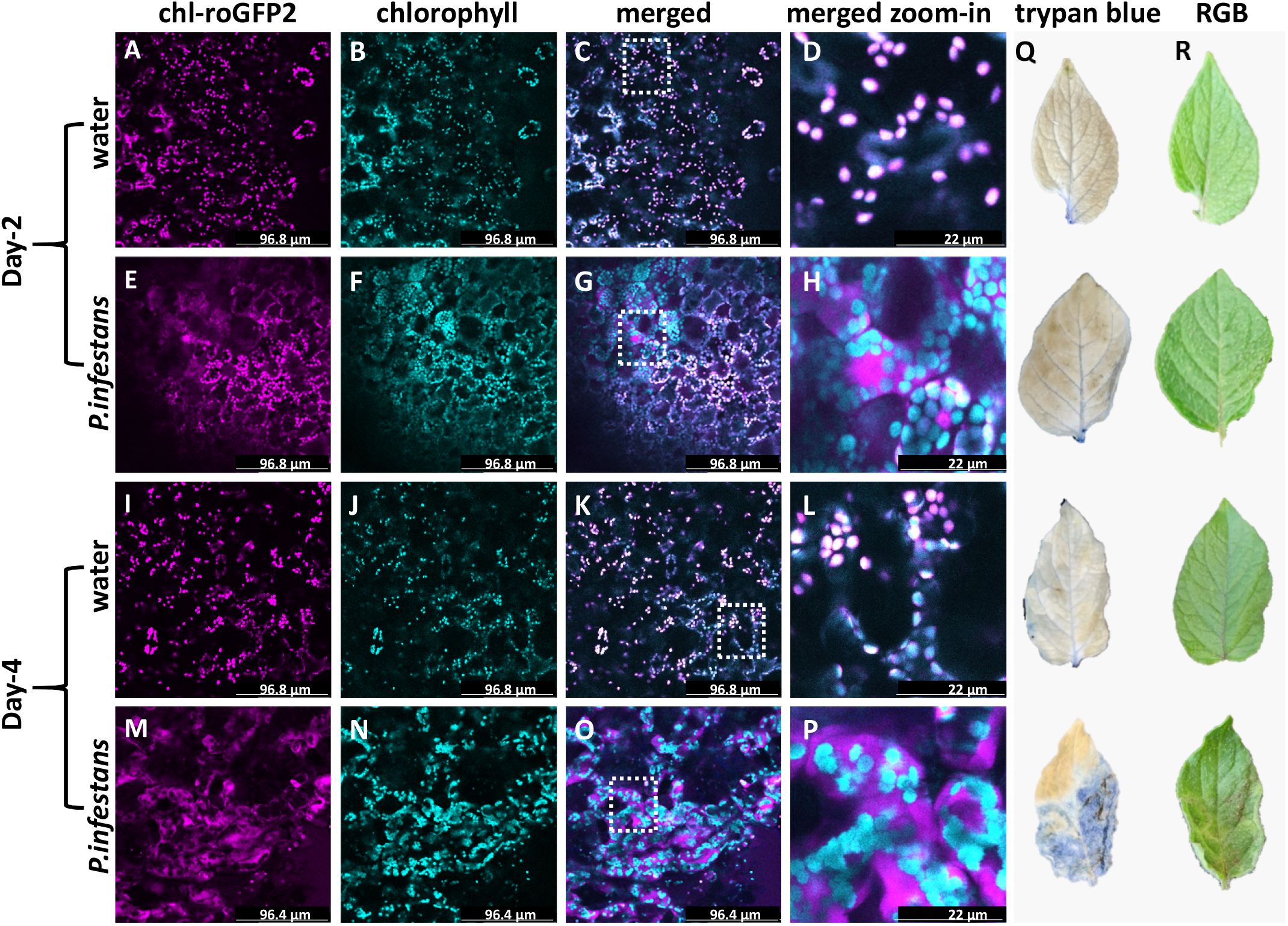
Confocal microscopy images of chl-roGFP2 in detached potato leaves infected with *P. infestans*. A, E, I&M) Chl-roGFP2 flouresence following excitation at 488 nm. B, F, J&N) Chlorophyll autoflouresence following excitation at 488 nm. C, G, K&O) Merged images of chl-roGFP2 and chlorophyll flouresence. White dashed square indicates the zoom-in zones. D, H, L&P). Merged magnified zone of chl-roGFP2 and chlorophyll. Q) Trypan blue staining to assess cell death in detached leaves from the 2 and 4 days post-inoculation (dpi). R) RGB images of detached leaves from the 2 and 4 dpi.

#### Redox imaging of dark-treated leaves detects the biotrophic stage of *P. infestans* infection

We have previously reported that light-to-dark transitions induced chl-roGFP2 oxidation in Arabidopsis plants, possibly reflecting the transmission of chloroplast-originating oxidative signals associated with the shutoff of photosynthetic enzymes (Haber et al., 2021). We therefore hypothesized that the mislocalized chl-roGFP2 probe observed in *P. infestans-* infected cells have lost its responsiveness to light-to-dark transitions. If true, analysis of redox images taken under light-to-dark transitions might identify the infected zones during the biotrophic phase. Indeed, chl-roGFP2 oxidation was observed in response to light-to-dark transition in control uninoculated leaves, whereas *P. infestans*-infected leaves displayed at 2 dpi distinct spots of reduced chl-roGFP2 oxidation surrounded by areas with higher chl-roGFP2 oxidation (D-R_405/465_; Fig. 3 & Supp. Fig. 5B &C). Notably, closer inspection of the localization of chl-roGFP in the reduced spots compared to the surrounding area associated with higher chl-roGFP2 oxidation verified that the reduced sites were associated with chl-roGFP2 mislocalization (Fig. 4 A-E). Particularly, 30% difference was found between the chl-roGFP2 oxidation level in areas exhibiting mislocalized chl-roGFP2 compared to the surrounding leaf tissue, demonstrating a deviation of 18 mV in E_GSH_ within the same leaf (Fig 4, D-E). In order to further demonstrate that the reduced spots observed in D-R_405/465_ images are associated with active infection, we infected specific sites on the leaves with droplets containing *P. Infestans*. In agreement with the spray inoculation results, D-R_405/465_ images at 2 dpi showed circular reduced spots in the infection locations, confirming that the reduced spots indicate infected areas (Supp. Fig. 6B).

**Figure 3:**
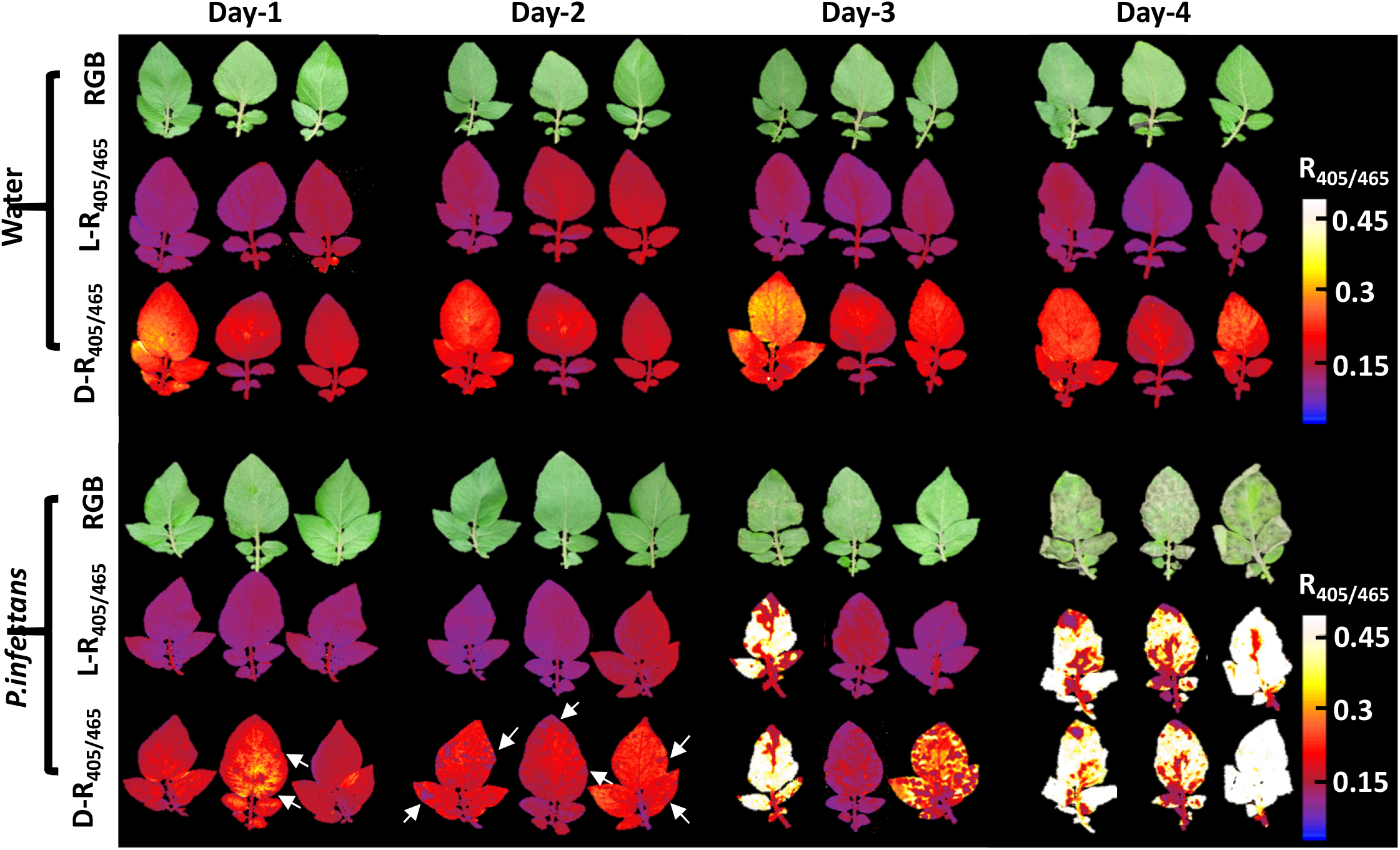
Redox imaging during light-to-dark transition of detached potato leaves infected with *P. infestans*. Detached leaves were sprayed with water or *P. Infestans*. Representative images show leaves in RGB, ratiometric imaging under light (L-R_405/465_) or light-to-dark transition (D-R_405/465_). White arrows indicate the spots exhibiting low chl-roGFP2 oxidation state*s*.

**Figure 4:**
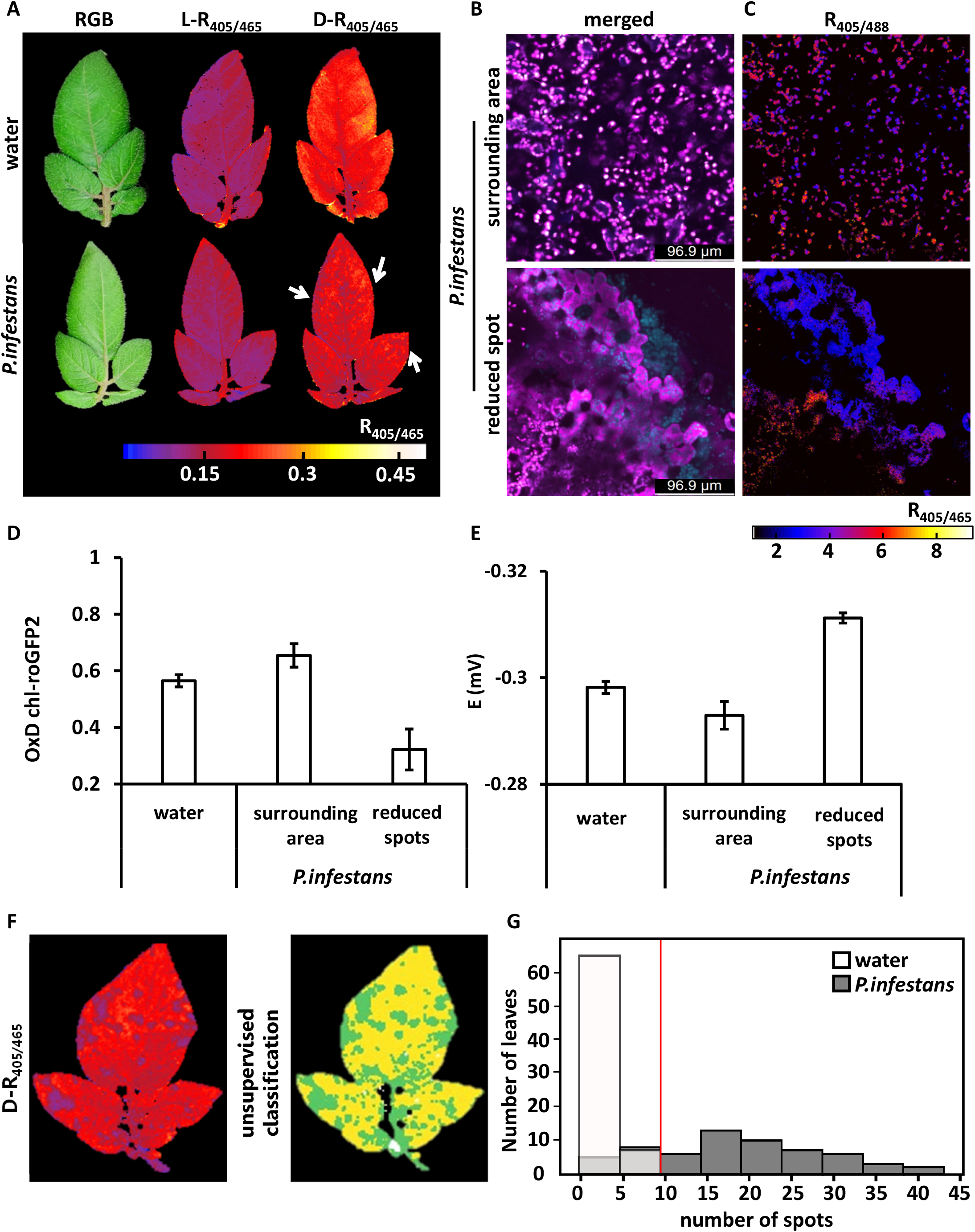
Chlorophyll fluorescence and ratiometric (D-R_405/465_) imaging of potato leaves inoculated with *P. infestans*. A) The maximum quantum yield of PSII (Fv/Fm), and non-photochemical fluorescence quenching (NPQ) images of fresh leaves on day 0. B) Images as in A plus ratiometric imaging of detached dark-adapted leaves (D-R_405/465_) on the second day post-infection (dpi). ΦPSII and NPQ were determined from the sixth pulse of actinic light. White arrows indicate spot locations. C) Comparison between Fv/Fm values obtained from spots exhibiting reduced chl-roGFP2 (R), the surrounding area (S) and water-treated leaves (W)as determined based on the D-R_405/465_ images in infected leaves. Values represent mean ± SE (*n*□= □18) of 3 spots from 6 different leaves. D, E) NPQ (D) and ΦPSII (E) induction in spots exhibiting reduced chl-roGFP2 spots, as determined from the D-R405/465 images *of P. infestans*-infected leaves. Values represent mean ± SE (*n*□= □18), composed of 3 spots from 6 different leaves.

To determine whether infected leaves which exhibit mislocalized roGFP2 areas are still responsive to redox alterations, we treated infected leaves at 2 dpi with DTT or H_2_O_2_ and followed the changes in R_405/465_ values. R_405/465_ values of 0.07 and 0.33 were measured for DTT and H_2_O_2_, respectively, demonstrating a roGFP2 dynamic range of 4.7 (Supp. Fig. 7B). A similar dynamic range was measured in non-infected leaves (Fig. 1, Hipsch et al., 2021), demonstrating the preservation of probe responsiveness to redox changes during infection.

The structured autofluorescence was assessed to verify that the observed changes in R_405/465_ are not affected by a possible alteration in autofluorescence in specific leaf areas in response to *P. infestans* infection by recording emissions at 435–485□nm following excitation at 405□nm (Fricker, 2016). As shown in Supp. Fig. 6&8, autofluorescence recorded on 2 dpi was minor compared to chl-roGFP2 signals. Accordingly, ratiometric analysis of fluorescence images that were quantitatively corrected for structural autofluorescence bleed-through into the biosensor channels revealed the same patterns of reduced spots during the biotrophic infection stage (Supp. Fig. 6&8). While leaf patches exhibiting increased autofluorescence were seen atthe necrotrophic stage (4 dpi), images corrected for this increase demonstrated higher oxidation state of the roGFP2 probe in these areas. These results indicate that the reduction and oxidation patterns on the potato leaves during *P. infestans* infection were not affected by the leaf autofluorescence.

#### Quantifying infection spots by machine learning-based image analysis

To obtain high-throughput *in-planta* detection of *P. infestans* in inoculated leaves we combined redox imaging with automated image analysis. An unsupervised machine learning (ML) classification algorithm was used to establish an automated pipeline to isolate patches from the leaf matrix (Fig. 4 F). the number of spots (i.e., infected zones) was counted using the scikit-image algorithm, while tiny (<1% of the total leaf area) and large (>30%, more than a third of the total leaf area) patches were removed (Van der Walt et al., 2014). As shown in Fig 4, F,G, testing of 72 non-infected vs. 60 infected leaves at 2 dpi resulted in a bimodal distribution with a clear cut-off threshold of more than 10 spots per leaf for infected leaves (Fig. 5 G). Though some infected leaves had fewer spots than this threshold, no non-infected leaves were classified as infected. These results demonstrate an unbiased quantification of the number of chl-roGFP2-marked spots generated during *P*.*infestans* infection and show how redox images and ML algorithms can be combined to automatically identify infected and non-infected leaves.

**Figure 5:**
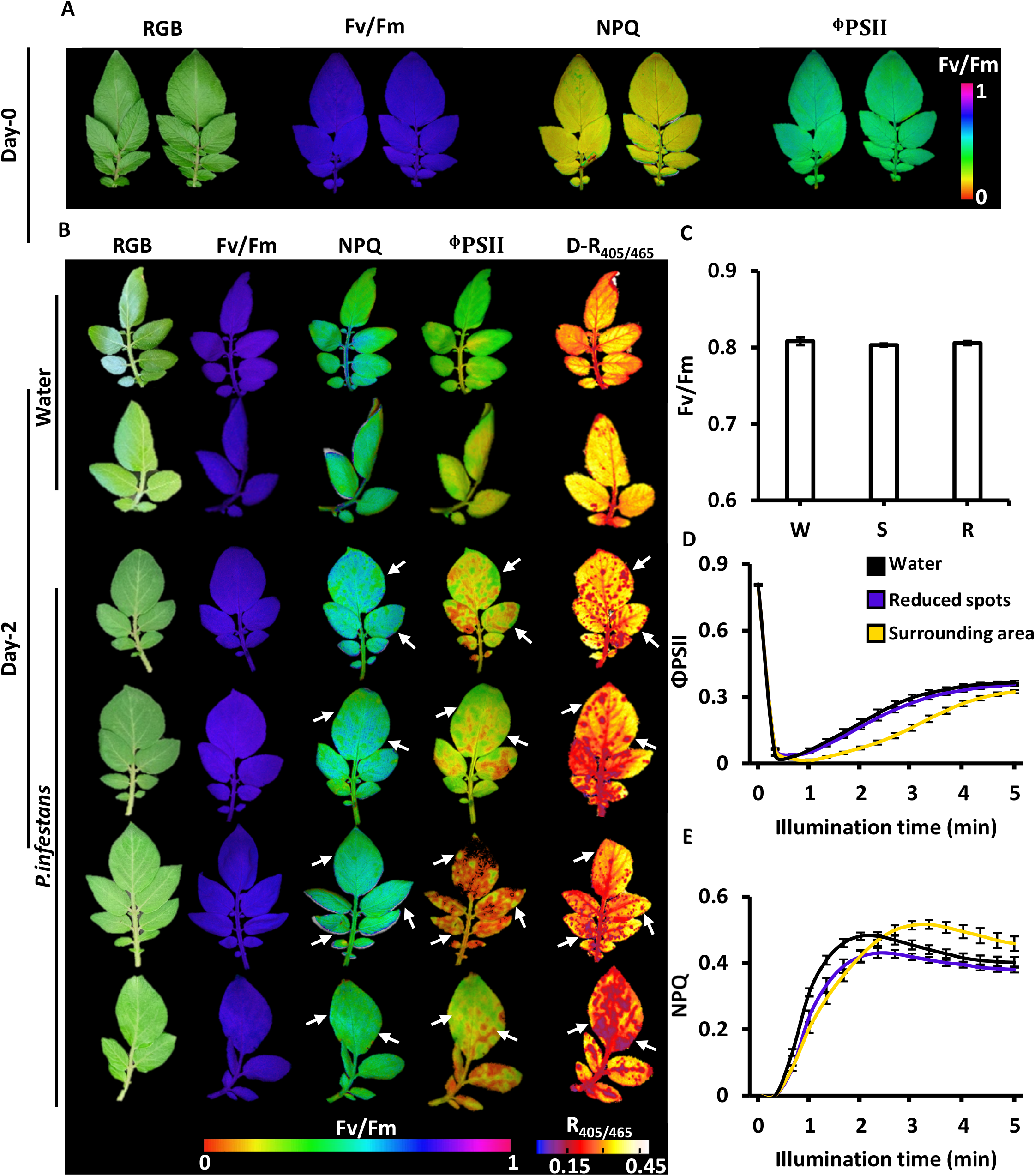
Redox imaging during light-to-dark transition identifies *P. infestans* infection zones in potato leaves on the second day post-inoculation. A) Representative images show *P. infestans-*infected leaves or control in RGB, ratiometric imaging (L and D-R_405/465_). White arrows indicate the spot locations. B) Confocal images showing the merged images of chl-roGFP2 and chlorophyll fluorescence (excitation: 488, 552 nm). C) Ratiometric analysis based on D-R_405/465_ images of different zones in spots exhibiting reduced chl-roGFP2 spot and the surrounding areas D&E) chl-roGFP2 oxidation degree (OxD, D) and GSH redox potential (E) in spots exhibiting reduced chl-toGFP2 in *P. infestans*-infected leaves in comparison to the surrounding area or water-treated leaves on the second day post-infection (DPI) (n=6). F) An example of unsupervised classification applied on a single D-R_405/485_ image of an infected leaf, 2 DPI. The unsupervised machine learning model was used to classify the leaf area into two areas: infected zones (green) and matrix (yellow). Spots were then recorded and counted per leaf in 72 control and 60 infected leaves, G) showing a clear cutoff of >10 spots per leaf (vertical red line) for infected leaves.

#### Mislocolized chl-roGFP2 in the early stages of *P. infestans* infections is associated with improved photosynthesis parameters

The efficiency of the photochemical electron transport chain in the infected zones as revealed by the redox imaging, was assessed on 2 dpi by measuring NPQ and quantum yield of PSII (ΦPSII). Comparing freshly detached leaves versus those kept for two days after detachment indicated an increase in NPQ and a slight decrease in ΦPSII values, independent of infection (Fig. 5 A, Supp. Fig. 9). Comparing infected and non-infected leaves resulted in similar Fv/Fm values in both groups, with no unique spots identified (as already indicated above in Fig. 1). Notably, spots exhibiting lower NPQ values of ∼0.42, surrounded by tissue with NPQ levels of ∼0.52, were noted in *P. infestans*-treated leaves. Similarly, the spots also exhibited higher ΦPSII values as compared to their surrounding tissue (0.27 vs. 0.13, respectively), suggesting improved photosynthesis efficiency due to infection (Björkman & Demmig, 1987; Skillman, 2008). Remarkably, a partial overlap was observed between the spots exhibiting low D-R_405/465_ and those showing altered NPQ and the ΦPSII, suggesting that mislocalization of chl-roGFP2 is associated with changes in photosynthetic efficiency.

#### Redox imaging distinguishes between infected and non-infected leaves at the whole-plant level

Following the detection of spatially resolved redox patterns induced by P. infestans in detached leaves, we assessed the potential to screen for infected leaves in intact potato plants grown in soil. As shown in Fig. 6 A, leaves displaying reduced spots, similar to those detected in detached leaves, were identified by whole-plant redox imaging of inoculated potato plants expressing chl-roGFP2. Suppression of photosynthesis is one of the earliest physiological responses to pathogen attack (de Torres Zabala et al., 2015). Thus, we compared the net photosynthetic CO_2_ assimilation in leaves exhibiting highly reduced chl-roGFP2 spots vs. leaves in the same plants with no reduced spots. As shown in Fig. 6 A, actively infected leaves, as evidenced by reduced spots, showed a lower carbon assimilation rate than uninfected leaves (Fig. 6 B&C).

**Figure 6:**
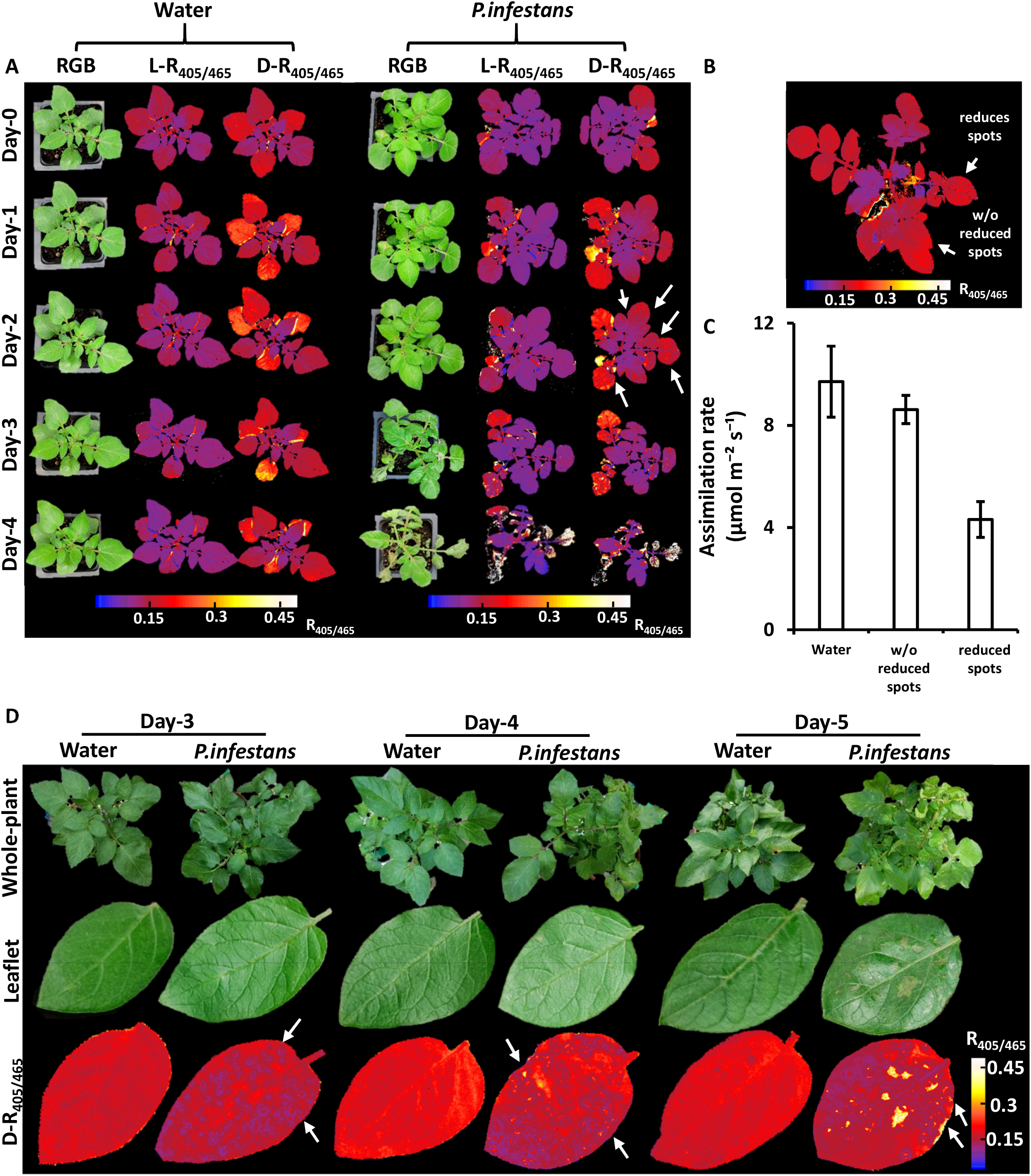
Redox imaging detected late blight on the whole-plant level and under natural greenhouse conditions. A). Representative images of plants infected with *P. infestans* or sprayed with water as control. RGB and ratiometric (L-R_405/465_ and D-R_405/465_) images are shown. White arrows indicate leaves exhibiting spots with reduced chl-roGFP2, a sign of active *P. infestans* infection. B) A representative image of the *P. infestans-*infected plant showing infected and uninfected leaves in the same plant. C) Carbon assimilation rates measured in potato plants infected with *P. infestans*, from the second day post-inoculation (dpi). The leaves were classified as non-infected (no reduced chl-roGFP2 spots detected) or infected (based on detected reduced chl-roGFP2 spots). Carbon assimilation rates were measured under the illumination of 400 photons m^−2^ s^−1^. Values represent mean ± SE (n□= □4-5). D) Potato plants grown in the greenhouse were infected with *P. infestans* or sprayed with water as control. Representative RGB and ratiometric images (D-R_405/465)_ taken of detached leaflets are shown. White arrows indicate signs of *P. infestans* infection.

Finally, we examined redox images derived from infected plants grown in a greenhouse (Fig 6, D). To this end, leaflets detached from infected and non-infected chl-roGFP2-expressing potato plants were imaged daily to resolve changes in the probe redox state. Notably, while visual symptoms associated with a shift to the necrotrophic phase were recognized at 5 dpi, reduced spots indicative of the biotrophic stage were already observed on 3 dpi. Inspection of the redox images showed a slightly different redox pattern than that obtained under environmentally controlled conditions. The spots had an outer ring-like pattern of highly reduced roGFP2 and a higher oxidation state in their center (Fig. 6, D). Taken together, by triggering chl-roGFP2 mislocalization outside the chloroplast, *P. infestans* infection generates a unique redox signature on the whole-plant level during light-to-dark transition under natural growth conditions.

### Discussion

Despite the 180 years that have passed since the Irish “Great Famine”, *P. infestans* still poses challenges to agriculture practices (Guenthner et al., 2001). Recent studies have pointed out the prominent role chloroplasts play in mediating plant-pathogen interactions through their generation of defense signals such as plant hormones and ROS (de Torres Zabala et al., 2015; Park et al., 2018; Serrano et al., 2016). In addition, chloroplast-targeted pathogen effectors were identified and shown to be involved in manipulating and altering chloroplast shape, formation and function (de Torres Zabala et al., 2015; Gao et al., 2020; Germain et al., 2018; Khoury et al., 2014; Petre et al., 2015; Savage et al., 2021; Xu et al., 2019; F. Yang et al., 2021).

The primary goal of this study was to examine the effect of *P. infestans* on the chloroplast *E*_*GSH*_. To achieve high spatiotemporal resolution, whole-plant fluorescence imaging was used to detect changes in chl-roGFP2, as done recently under various abiotic stress conditions (Hipsch et al., 2021a). Unexpectedly, chl-roGFP2 was found mislocalized outside the chloroplast during the presymptomatic biotrophic stage of the infection, before visual symptoms were apparent (Fig. 2&3). While chl-roGFP2 mislocalization precluded the monitoring of chl-*E*_*GSH*,_ it suggested the impairment of the chloroplast protein import system during the biotrophic stage of late blight disease and allowed the development of a new approach for nondestructive monitoring of the early presymptomatic state of the disease at the whole-plant level.

While chloroplasts can synthesize proteins encoded in the chloroplast genome, most chloroplast proteins are nuclear-encoded, and their import from the cytosol is mediated by two complexes in the outer and inner envelope membranes, the TOC and TIC, respectively (Paila et al., 2015; X. Yang et al., 2019; Yu et al., 2012). Tight regulation of the import process is critical in shaping the chloroplast proteome during plant development and under abiotic stress conditions (Ling et al., 2012; Ling & Jarvis, 2015). Specifically, TOC inhibition under abiotic stress conditions, meditated by the envelope-localized ubiquitin E3 ligase SUPPRESSOR of PPI1 LOCUS1, was shown as a strategy to mitigate harmful conditions by limiting the import of photosynthetic apparatus components, reducing the production of photosynthesis-derived ROS (Ling & Jarvis, 2015). Similarly, the mislocalization of chl-roGFP2 outside the chloroplast detected in response to *P. infestans* infection likely reflects the mislocalization of natural chloroplast proteins. In adition to reducing ROS production, mislocalization can act as a host immunity response by inhibiting the import of chloroplast-targeted pathogen effectors that manipulate chloroplast metabolism. Interestingly, quantitative analysis of reduced spots on the whole-leaf level, revealed approximately 25 patches with mislocalized chl-roGFP2 per leaf at the biotrophic infection stage, under the infection conditions applied in this study, in which many of then did not develop into necrotic patches (Fig. 5). The development of considerably fewer necrotic patches at the necrotrophic stage suggests that the elicited immune response prevented disease progression. Accordingly, crop losses by plant pathogens may be curbed by applying rational engineering strategies to manipulate the protein import system.

Leaf-level interactions between plants and pathogens are governed by unique spatiotemporal dynamics. The invisibility of the biotrophic stage makes it challenging to differentiate between cells under active infection or immune response and the surrounding non-infected cells. Accordingly, most studies exploring, for example, gene expression patterns, average the entire leaf area, ignoring the spatiotemporal heterogeneity in the infection state. One of the most important observations in this study was that infection spots were spatially resolved due to the mislocalization of chl-roGFP2. This was achieved thanks to the loss of responsiveness of chl-roGFP2 to light-dark transition in the mislocalized areas, leaving leaf patches with a lower chl-roGFP2 oxidation state (Fig. 3). By exploiting the ability to nondestructively and spatially resolve infection spots, redox imaging can serve as an ideal tool for investigation of the molecular changes in tissues at different stages of infection. For example, the combination of redox imaging and spatial transcriptomics or mass spectrometry imaging (Dong et al., 2016; Giacomello et al., 2017; Petras et al., 2017; Schleyer et al., 2019; Solanki et al., 2020) has the potential to precisely tap into the genetic and metabolic processes underlying disease onset.

Potato plants live in highly inconsistent environments, and continuous exposure to minor stress levels interrupts plant homeostasis and results in constant energy loss due to resource diversion toward the activation of defense mechanisms (Fahad et al., 2017; Gold, 2021; Hipsch et al., 2021a; Zhu, 2016). Various technologies have been developed for the early detection of plant pathogens, such as nucleic acid-based methods and multispectral imaging (Baldeck & Asner, 2014; Gold et al., 2020; Khan et al., 2017; Lees et al., 2012; Zhao et al., 2016). This work is a critical step toward developing new quantitative tools for the detection of late blight disease in crop plants and controlling and limiting disease spread by utilizing genetically-encoded biosensors. Coupled with a portable fluorescence detection system and automatic ML-based image analysis, integration of this new methodology in agricultural fields has the potential to dramatically improve high-throughput phenotyping capabilities in plant breeding programs and optimize the management of one of the most devastating diseases in potato crops.

### Materials and Methods

#### Plant material and production of transgenic plants

Wild-type and chl-roGFP2-expressing potato (*Solanum tuberosum)* plants were planted in soil in 27 × 54 cm pots and placed in a controlled-environment growth chamber (60–70% relative humidity and ambient CO_2_ and under 14-h light [120 µmol m^−2^ s^−1^]/10-h dark per 24 h cycle. Plants were vegetatively propagated from cuttings or tubers. Unless otherwise stated, experiments were performed on 3–to 4-week-old plants in a FytoScope FS-RI 1600 plant growth chamber (Photon Systems Instruments). Plants were relocated to the chamber several days before the experiments started to allow acclimation to the chamber environment.

For the greenhouse experiment (conducted in Rehovot, Israel, in February 2022) 4-week-plants were grown from cuttings in 4 L pots filled with sandy soil medium. Plants were spray-inoculated with sporangial suspension (1×10^5^ sporangia per ml) of *P. infestans* (genotype 23_A1), wrapped with plastic bags and incubated in a growth chamber in the dark at 18°C. Bags were removed after 18h and plants were maintained in the growth chamber under photoperiod conditions. Plants sprayed with DW served as control. At various time intervals after inoculation, randomly selected leaflets were detached from the 4th-6th node from the top of the plants and placed in a closed humidified and illuminated chamber for the imaging processes (up to 60 min).

#### Pathogen and inoculation

Isolate 164 *Phytophthora infestans (P. infestans)* (genotype 23_A1, resistant to mefenoxam) was used in all experiments. It was collected in March 2016 from a potato field at Nirim, Western Negev, Israel (Cohen, 2020). It was propagated in a growth chamber at 18°C by repeated inoculations of detached potato leaves. For inoculation, freshly produced sporangia were collected at 5-7 dpi from infected leaves by washing with distilled water (DW) into a beaker kept on ice (4°). Sporangial concentration was adjusted to 1×10^5^ sporangia per 1 ml with the aid of a hemocytometer. Detached leaf assays were done with leaves from the 2th and 3th node from the top of the plants. Leaves were sprayed on the abaxial side with sporangial suspension, placed on a sterile moist filter paper Whatman No 1. inside petri dishes and kept at 18°C in the dark for 24 h (Cohen et al., 1991). Leaves were then incubated in a growth chamber as above. Control leaves were sprayed with sterile DW. Whole-plant inoculations were performed with 3-4-week-old plants. They were sprayed with sporangial suspension of *P. infestans*, covered with plastic bags for 18-24 h and maintained in dark conditions.

#### Confocal microscopy

Images were acquired with a Leica SP8 Lightning Confocal system (Leica Microsystems) and LAS X Life Science Software equipped with a HCX APO U-V-I 40×/0.75 DRY UV objective. Images were a cquired at 4096□× □4096-pixel resolution, with 507–534 nm emission bandpass and photo multiplier tube (PMT) gain of −630(V) following excitation at 488 nm or 405 nm for chl-roGFP2 fluorescence detection. For the 488 and 405 nm excitation, 15% and 30% of the maximum laser intensity were used, respectively. The 652–692 nm emission bandpass and PMT gain of −571.4(V) following excitation at 488 nm were used to image chlorophyll fluorescence. Autofluorescence following excitation at 405 nm was recorded at 431–469□nm. Merged images were generated using LAS X software. Subtraction of mean autofluorescence values from 405 nm images of a user-defined region of interest (ROI) that did not include chloroplasts was performed to remove background signals. The ratiometric images were generated by dividing, pixel-by-pixel, the 405 nm by the 465 nm signal, and displaying the result in false colors using Matlab.

#### Plant cell death staining

To visualize cell death in detached leaves, control and infected leaflets were detached 2 and 4 dpi and placed in trypan blue solution for 45 min, agitated in a lab rotator, and washed with absolute ethanol three times. Chlorophyll was removed from the leaflets by soaking them in absolute ethanol for two consecutive days.

#### *Phytophthora infestans* staining

*P. infestans* mycelia and sporangia were stained with 0.01% calcofluor white (18909-100ML-F, Sigma-Aldrich) in DW. Control and infected leaflets were detached, placed in calcofluor white solution for 45 min, agitated on a lab rotator, and washed twice with phosphate-buffered saline (PBS). Fluorescence was detected using an Advanced Molecular Imager HT (Spectral Ami-HT, Spectral Instruments Imaging, LLC., USA). Excitation was performed with 405 nm□± □10 light sources and a 490 nm □± □10 emission filter was used. Images were pre-processed using a custom-written Matlab script. Representative images were generated using the Aura program.

#### Chlorophyll fluorescence measurements

Chlorophyll fluorescence was measured using a Walz PAM IMAGING PAM M-series IMAG-K7 (MAXI) fluorometer. In all the experiments, the leaves were dark-adapted for 20 min. The maximum quantum yield of PSII (Fv/Fm) was determined by *Fv* / *Fm* = (*Fm* – *F0*) / Fm. Nonphotochemical quenching (NPQ) each at saturating pulse was determined by the equation 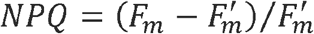 and the effective PS II quantum yield (ΦPSII) was determined by the equation of ΦPSII= (*Fm* - *F*)/*Fm*. The actinic illumination was set to 334 μmol photons m^−2^ s^−1^ and saturating pulses were applied at 20 s intervals for 5 min. Images were analyzed with ImagingWinGigE V2.56p software.

#### Gas exchange measurements

Leaf CO_2_ assimilation rates were measured using the portable gas analyzer Li-Cor-6800 gas exchange (LICOR, Lincoln, NE, USA). The leaf chamber maintained a constant CO_2_ level of 400 ppm. The temperature and humidity were set at 18°C and 70%, respectively.

#### Chl-roGFP2 fluorescence measurements and image analysis

Whole-plant chl-roGFP2 fluorescence was detected using an Advanced Molecular Imager HT (Spectral Ami-HT, Spectral Instruments Imaging, LLC., USA), and images were acquired using the AMIview software. For chl-roGFP2 fluorescence detection, excitation was performed with 405 nm±10 or 465 nm±10 LED light sources and a 515 nm±10 emission filter was used. For chlorophyll detection, a 405 nm±10 LED light source and a 670 nm±10 emission filter were used. All images were captured under the following settings: exposure time = 1 s, pixel binning = 2, field view (FOV) = 25 cm, FOV for detached leaves =15cm and LED excitation power 40% and 60%, for 405 nm and 465 nm excitations, respectively. Excitation power for chlorophyll detection was 5%. Chlorophyll autofluorescence was measured to generate a chlorophyll mask, which was then used to select pixels that returned a positive chlorophyll fluorescence signal. Only those pixels were subsequently considered for the roGFP analysis. The average signal of WT plants without chl-roGFP2 was determined and used for background correction of the fluorescence signals of chl-roGFP2 plants. For chl-roGFP2 autofluorescence correction (Supp. Fig. 7), excitation was performed with 405 nm□± □10 LED and a 448 nm□± □10 emission filter was used; the excitation power was 60%. Autofluorescence measured from WT plants was used to assess the factor between the autofluorescence measured at 488 nm and the bleed-through to the roGFP emission channels. This factor was used to generate a scaled version of the autofluorescence, which was subtracted from the original roGFP images. Ratiometric images were created by dividing, pixel-by-pixel, the corrected 405 nm image by the corrected 465 nm image and displaying the result in false colors. Images were pre-processed using a custom-written Matlab script.

#### roGFP2 probe calibration

For calibration of the probe response, detached, fully expanded leaves were immersed in 1000 mM H_2_O_2_ or 100 mM DTT, and ratiometric images for fully oxidized and fully reduced states were then acquired. roGFP2 OxD (the relative quantity of oxidized roGFP proteins) was calculated for individual plants based on the whole-plant fluorescence signal, according to equation (1) (Meyer et al., 2007).

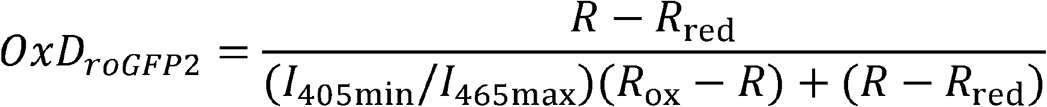

where R represents the 405/465 fluorescence ratio at the indicated time and treatment, R_red_ represents the 405/465 fluorescence ratio under fully reduced conditions, R_ox_-represents the 405/465 fluorescence ratio under fully oxidized conditions, I465_ox_- the fluorescence e mitted at 515 nm when excited at 465 nm under fully oxidized conditions and I480_red_- the fluorescence emitted at 515 nm when excited at 465nm under fully reduced conditions. E_GSH_ values were calculated according to Schwarzlander et al. (2008).

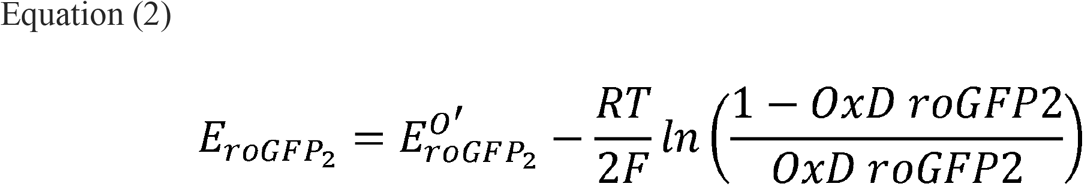

where 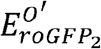 is the mid-point potential for the roGFP2, R is the gas constant (8.315 J K^−1^mol^−1^), T is the absolute temperature (298.15 K), and F is the Faraday constant (9.648 10^4^ C mol^−1^).

#### Machine learning-based image analysis

The following procedure was used to automatically identify images of infected leaves, distinguishing them from those of non-infected leaves using ML algorithms. First, each image was smoothed using a 3×3 pixel running window through the use of the scipy.ndimage.filters.generic_filter algorithm (Virtanen et al., 2020). Such a smoothing was necessary to remove small artifacts in the image, which sometimes showed random dots on and around the leaves. Then, the k-means clustering algorithm was applied to the cleaned image to classify the image into three clusters (Pedregosa et al., 2011). We used this procedure with different clusters, manually inspecting the images to see the results. We found that three is the optimum number of clusters that enables distinguishing the infected zones in the leaf, classifying the image into background area (class 1), leaf matrix area (class 2; non-infected zone), and infected area (class 3). The background area (class 1) was then removed, and the two remaining areas were segmented using the scikit-image algorithm (der Walt et al., 2014). By using the measure.label function that collects pixels with identical values, we were able to remove tiny dots (<4 pixels), which were obviously part of an artifact of the unsupervised classification k-means clustering algorithm. After normalizing each spot area (class 3) to the total leaf area (class 2 + class 3) (Barbedo, 2016), we further removed tiny spots (<1% of the total area of the leaf) and big spot areas (>33% of the leaf area), which were assumed to be part of the leaf matrix and not an infected area due to their large size. Finally, the number of remaining spots (class 3) was counted per leaf using the Scikit-image package (der Walt et al., 2014), and the distribution of the leaves by their number of spots was plotted as a histogram using the Matplotlib 2D graphic environment in Python (Hunter, 2007). Since images of infected leaves showed a spot-like pattern, we expected that the machine would classify images with leaves with the highest number of spots as infected. To obtain the cutoff threshold that distinguishes between infected and non-infected leaves, the mean of the means of the number of spots of 72 non-infected leaves and 60 infected leaves was calculated. Leaves with a higher number of identified spots than the cutoff threshold would be classified as infected.

