## Supplementary Figures for "Early detection of late blight in potato by whole-plant redox imaging"

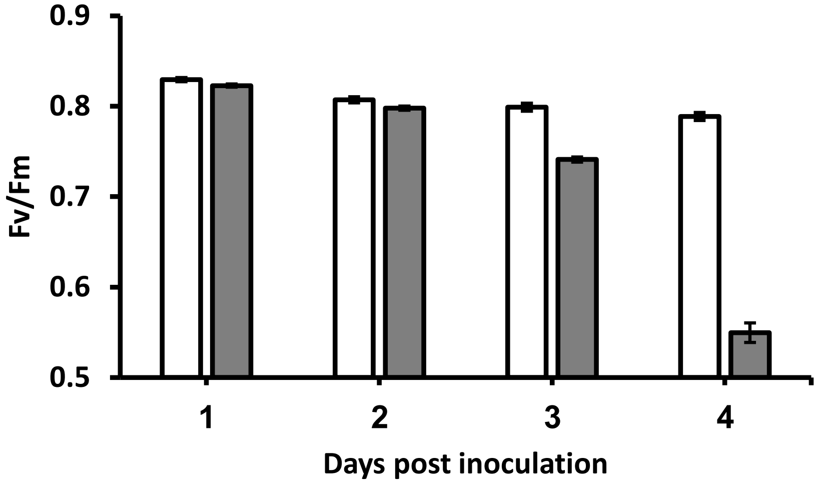


**Supplementary Figure S1: Maximum quantum yield of PSII (Fv/Fm) of potato leaves inculcated with water or *P.infestans.*** Fv/Fm values were derived from chlorophyll fluorescence imaging analysis and represent the means of 20 plants ± SE. The Fv/Fm analysis was carried out by randomly choosing eight circular areas of interest (AOI) in each leaf area.

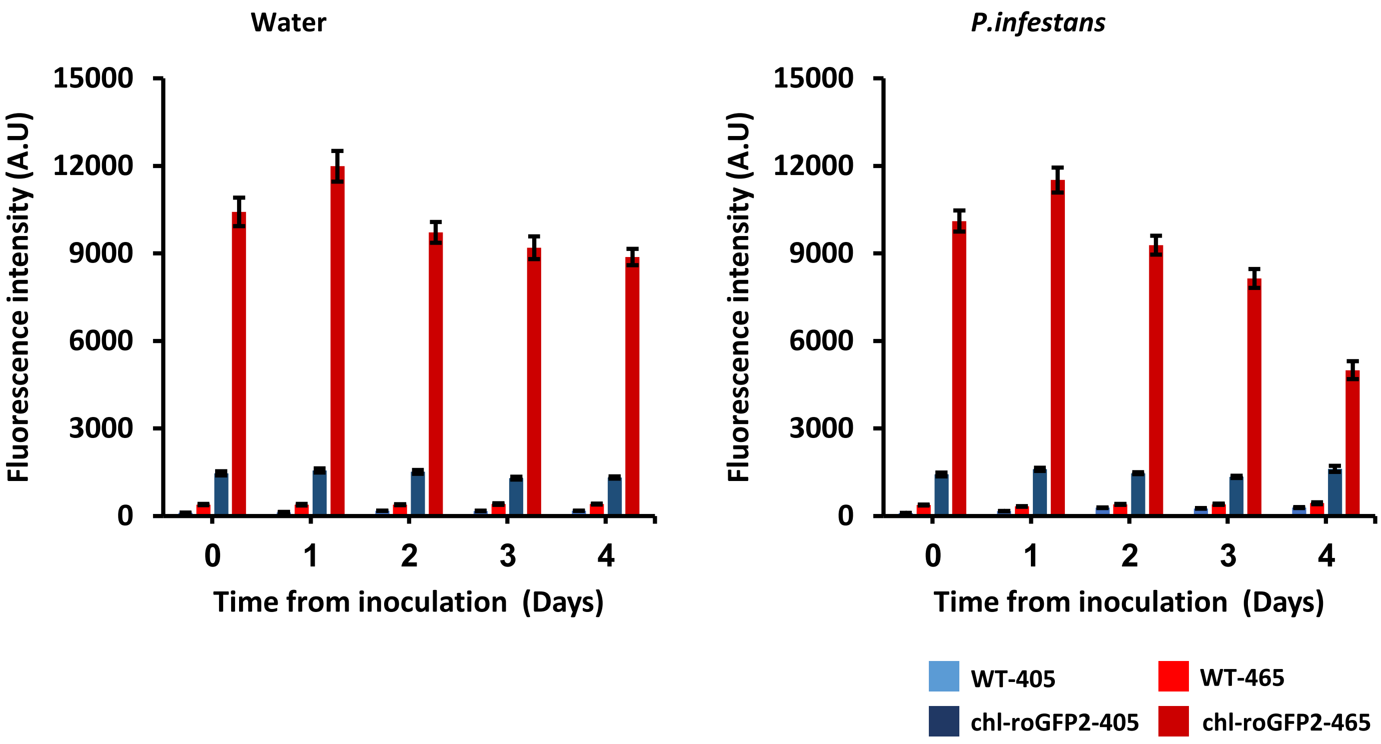


**Supplementary Figure S2: Fluorescence intensity values of chl-roGFP2 and WT leaves treated with water or *P.infestans* spores.** Raw fluorescence intensities values of the images shown in Fig. 1 are presented. Emission intensity values were recorded at 515 nm, following excitation at 405 or 465 nm. Values represent the means of 18-20 leaves ± SE for chl-roGFP2 plants and 5 leaves ± SE for WT.

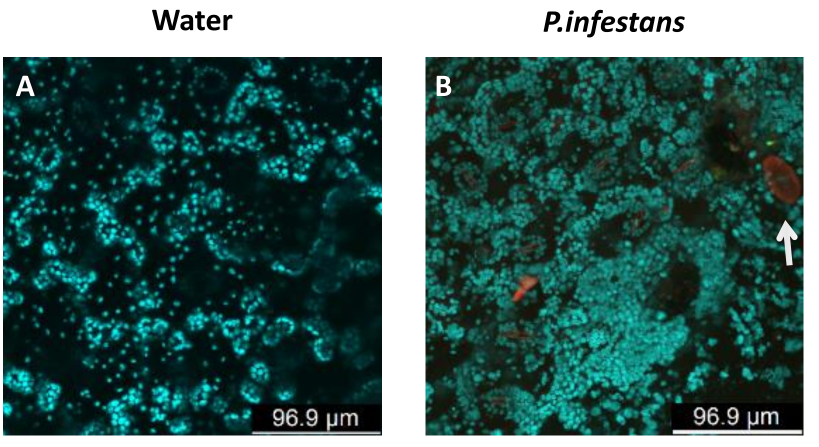


**Supplementary Figure S3: Chloroplast imaging of Potato plants treated with water or *P.infestans* spores on the second-day post-inoculation.** Confocal images of chlorophyll autofluorescence of WT leaves sprayed with water or *P.infestans* spores are shown. Chlorophyll autofluorescence was recorded at 488nm excitation. The white arrow indicates *P.infestans* spore.

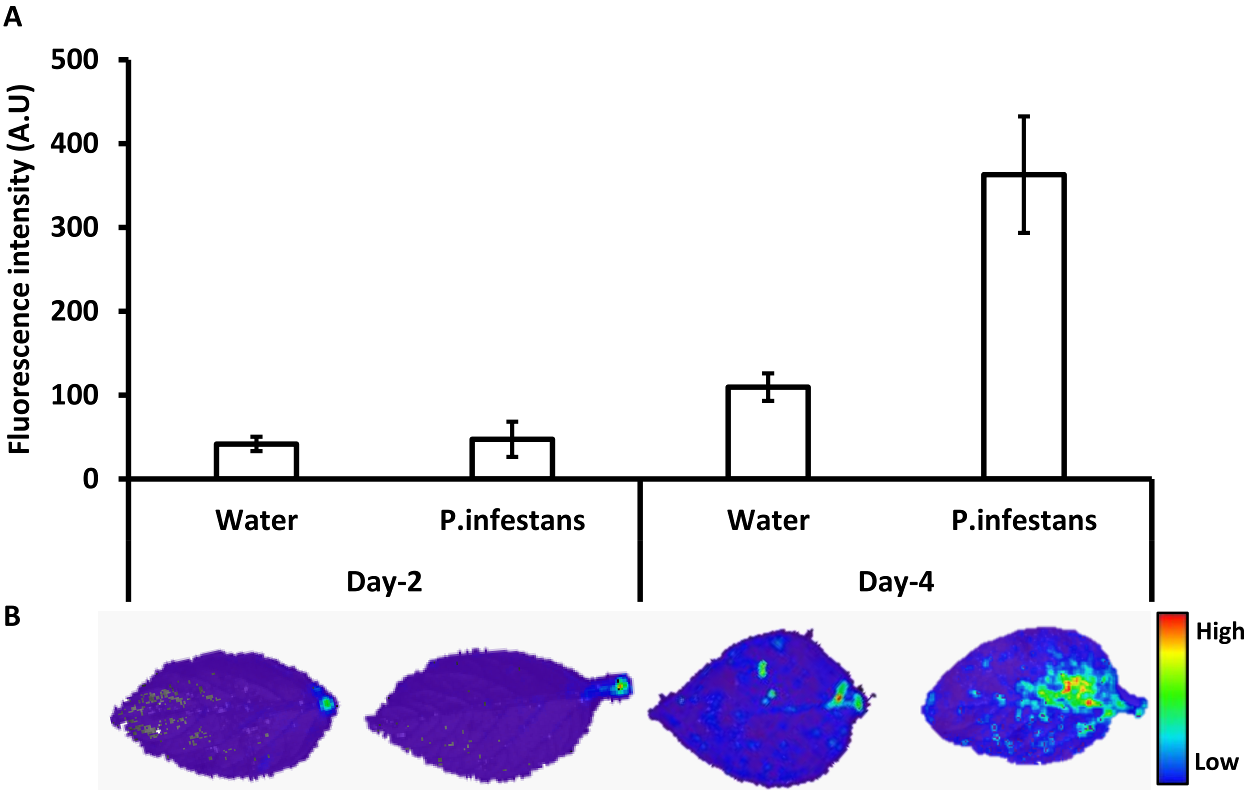


**Supplementary Figure S4: Calcofluor white imaging of uninfected and infected leaflets.** A) Quantification of the fluorescence intensities derived from detached leaflets stained with Calcofluor white on the 2 and the 4 dpi. Values represent mean ± SE ($n$ = 6-10). B) Fluorescence imaging of representative leaflets.

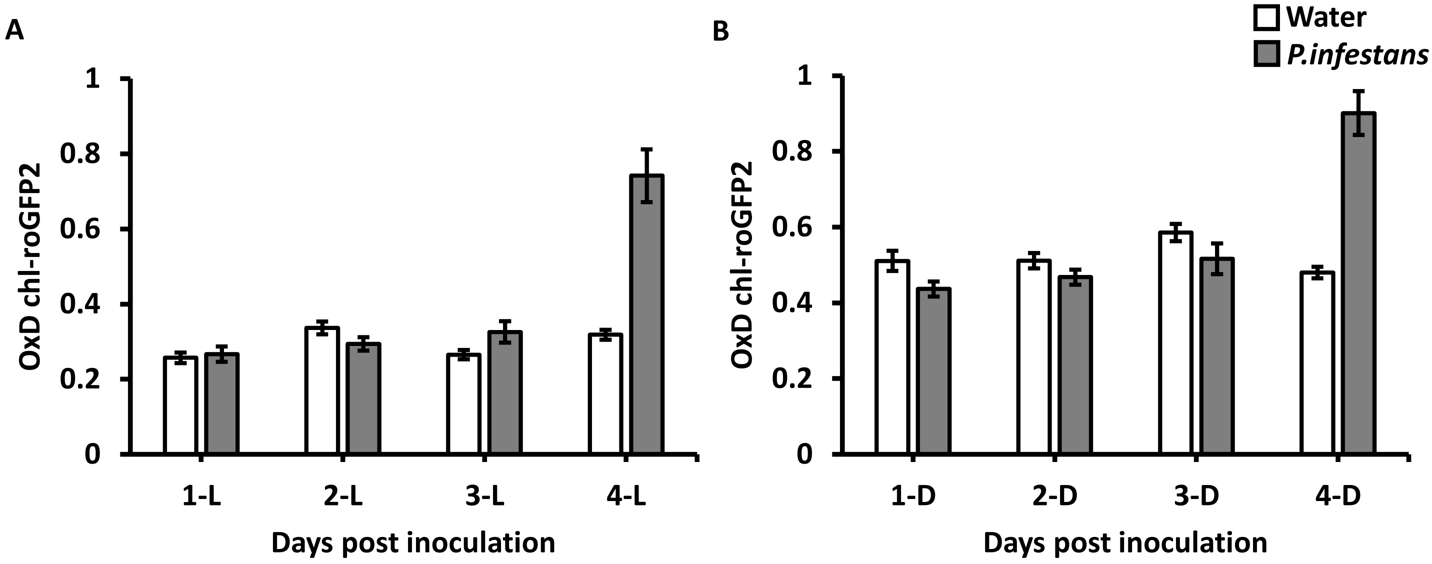


**Supplementary Figure S5: Oxidation degree measurements during the light and light-to-dark transition of detached potato leaves infected with water or *P. infestans*.** A) chl-roGFP2 degree of oxidation (OxD) in detached leaves inoculated with *P. infestans* or treated with water as measured in light-adapted leaves (B) or during the light-to-dark transition. Data represent mean ± SE (n = 20).

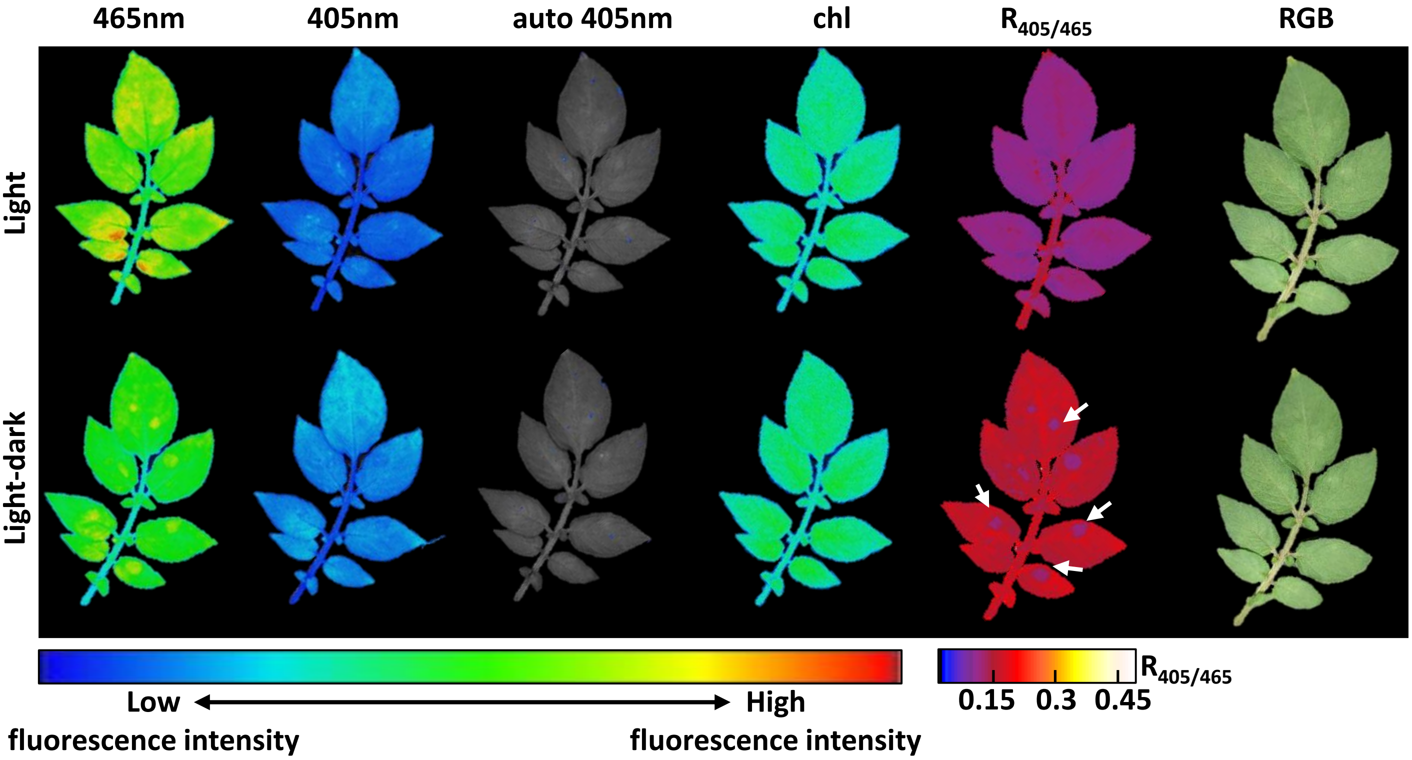


**Supplementary Figure S6. Imaging of leaves infected with droplets contain *P. infestans* on the second-day post-inoculation.** Detached leaf were inoculated with *P.infestans* droplets. Images of chl-roGFP2 fluorescence following excitation at 405 nm and 465 nm, autofluorescence following excitation at 405 (emission at 448nm), chlorophyll autofluorescence, autofluorescence-corrected ratiometric analysis and RGB images are shown. Images were taken from light-adapted leaves (Light) and during light to dark transition (Light-dark). White arrows indicate the locations of the droplets containing *P.infestans* in the D-$R_{405/465}$ image.

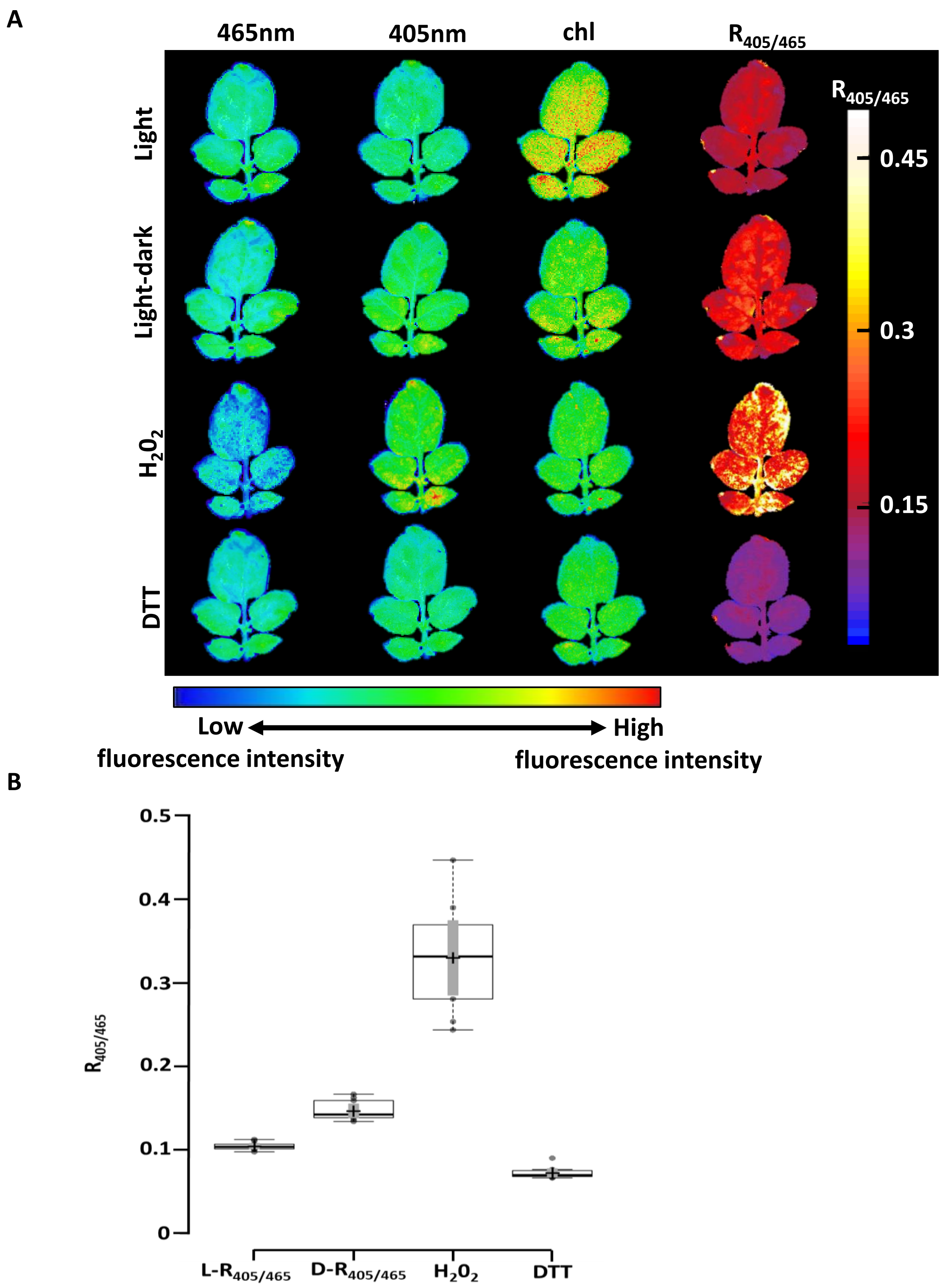


**Supplementary Figure S7: The responsiveness of infected chl-roGFP2 leaves to altered redox conditions.** A) Redox images of representative infected chl-roGFP2 expressing leaf on 2 dpi during adaption to light (Light), light to dark transition (Light-dark) and under fully oxidized (1,000 mM $H_2O_2$) and fully reduced (100 mM DTT) conditions. Images of detached leaves were digitally combined for comparison. B) Ratiometric analysis of leaf discs derived from infected chl-roGFP2 leaves in 2 dpi exposed to the conditions mentioned in A. Centerlines show the medians; box limits indicate the 25th and 75th percentiles as determined by R software; whiskers extend 1.5 times the interquartile range from the 25th and 75th percentiles, dots represent outliers; crosses represent sample means; bars indicate 95% confidence intervals of the means; width of the boxes is proportional to the square root of the sample size; data points are plotted as open circles (n = 10).

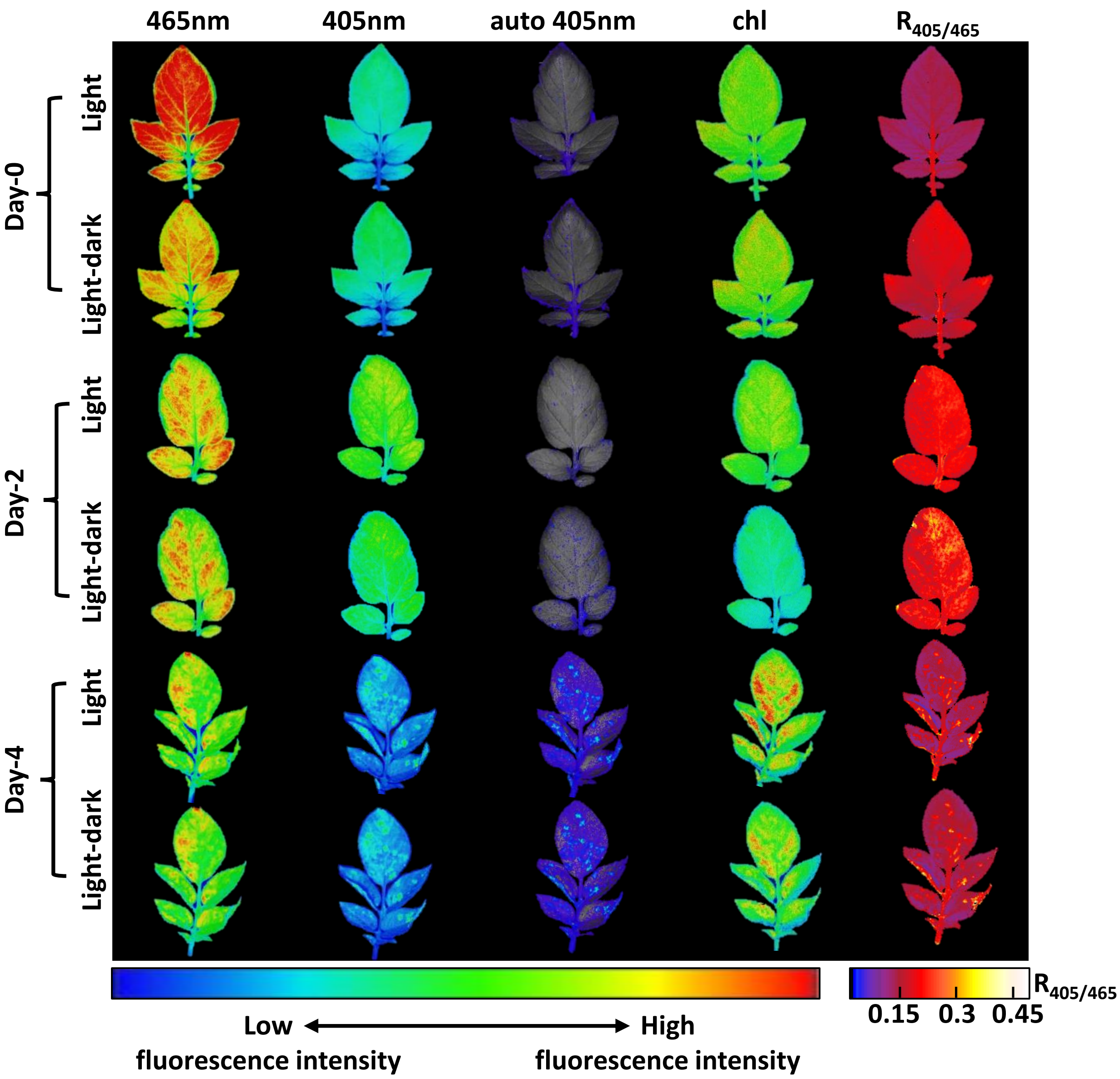


**Supplementary Figure S8: Assessment of structural autofluorescence background in chl-roGFP2 leaves inoculated with *P .infestans* inoculation.** Images of chl-roGFP2 fluorescence, autofluorescence following excitation at 405 (emission at 448nm), chlorophyll autofluorescence, autofluorescence-corrected ratiometric analysis and RGB images are shown. Images were taken on the 2 and 4 dpi from light-adapted (Light) leaves and during light to dark transition (Light-dark).

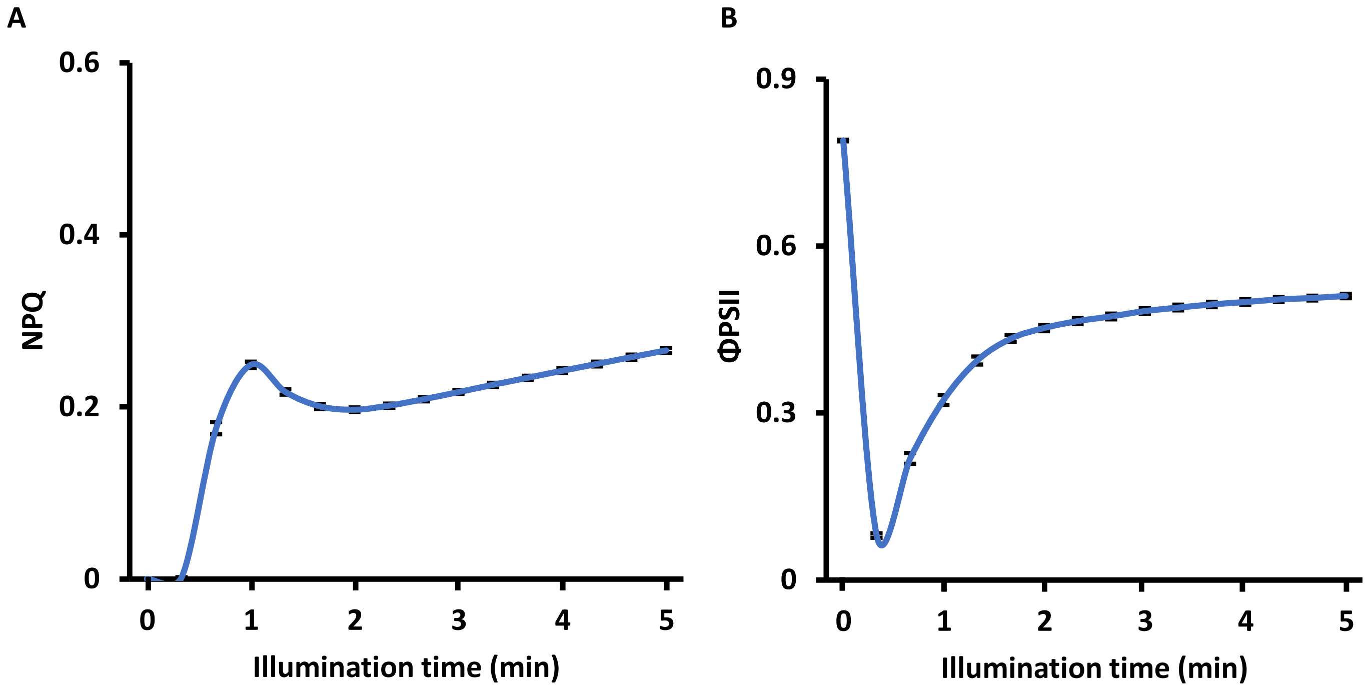


**Supplementary Figure S9: NPQ and ΦPSII values derived from freshly detached leaves.** NPQ (A) and ΦPSII (B) induction measurements during 5 min of illumination with actinic light intensity of 336 μmol photons $m^{-2}$ $s^{-1}$. The data represent the mean of 16 different leaves ± SE. In each leaf, values were collected from three randomly chosen circles.
